# Strong amplification of quantitative genetic variation under a balance between mutation and fluctuating stabilizing selection

**DOI:** 10.1101/2025.02.22.639683

**Authors:** Jason Bertram, Zahra Shafiei

## Abstract

The observation of high heritability in most quantitative traits has been a long-standing puzzle. There is a general consensus that simple models of quantitative genetic variation, which are mostly founded on the assumption of mutation-selection balance in a constant environment, have failed to explain high heritability. To make matters worse, the reasons for failure are unknown, leaving little to guide future model developments. Here we revisit this puzzle by taking the canonical Latter-Bulmer model and relaxing the assumption of perfect environmental stasis. Instead we assume that the trait optimum changes slowly but steadily in a random walk (specifically, an Ornstein-Uhlenbeck process), similar to standard models used for phylogenetic comparative methods. We show that our model behaves qualitatively differently to its stationary optimum counterpart even though the optimum only changes slowly. This is the result of a feedback between the adaptation rate and selection coefficient fluctuations. The heritability predictions resulting from this feedback are more consistent with observations and also less sensitive to evolutionary parameters than the classical LB model. We derive a simple formula to predict genetic variation under random walk optimum fluctuations which helps to explain some of our counter-intuitive results and which should be useful for understanding the potential influence of fluctuations in future work. Since the feedback driving our results should also occur in more complex models e.g. with multiple traits, we argue that environmental change has been an essential biological ingredient missing in previous mutation-selection balance models of quantitative trait heritability.

## 1 Introduction

Quantitative traits exhibit abundant genetic variation even when under strong selection, with narrow-sense heritability typically being in the range *h*^2^ = 0.1 − 0.6 [31, 21]. This is surprising because selection usually reduces genetic variation. We therefore expect to see abundant variation or strong selection, not both [38].

High variation despite strong selection can be explained by mutations if the influx of new variation is sufficiently large — the mutation-selection balance (MSB) hypothesis. To explore and test the MSB hypothesis, quantitative genetic models have been developed to predict variation under MSB. To date these models have failed to convincingly reconcile high heritability and strong selection because the only way to have both is to assume that the trait’s mutational target is huge [36, 21, 18, 39, 32]. Multiple lines of evidence support high polygenicity in many traits [32], including quantitative trait loci experiments [17], evolve and resequence experiments [23, 1] and genome-wide association studies. However, polygenicity appears to vary considerably between traits, and it remains unclear just how large we can safely assume mutational targets to be [32].

What is clear is that the demands of MSB models are severe. The canonical MSB models are the Latter-Bulmer (LB) [30, 7], and “house of cards” models [36]. Both models assume a single trait under direct stabilizing selection with a static optimum, and make identical heritability predictions. In *D. Melanogaster*, these models require ∼ 1% of the entire euchromatic genome to contribute to each trait [18] (also see Sec. 3.4). Moreover, predicted heritability is sensitive to the model parameters (mutation rate, intensity of selection and number of loci) [11]. Therefore, to be consistent with observations, canonical MSB models require tight constraints on evolutionary parameters including consistently large mutational target sizes [32, 11].

Here we show that the vanilla Latter-Bulmer model is much better at predicting realistic heritabilities if we relax the unrealistic assumption of perfect environmental stasis. Environmental fluctuations have long been recognized as an important determinant of genetic diversity [24, 5], and are ubiquitous in nature [25]. There is a sizeable literature on environmental fluctuations in genetics, but with an overwhelming focus on fluctuations as a source of balancing selection [6, 40, 42, 16]. Our understanding of mutation-selection-balance in a fluctuating environment is rudimentary in comparison.

Studies of single-locus models [22, 33] have found that selection coefficient fluctuations *reduce* heterozygosity [34] and produce an excess of derived major alleles [19] compared to the neutral case. Reduced variation is analogous to the loss of variation under stronger random genetic drift. However, these studies do not consider fluctuations a context of purifying selection on new mutations, which is an essential feature of stabilizing selection.

Studies of quantitative trait evolution in a changing environment have demonstrated substantially elevated genetic variation under directional [27, 9, 10], cyclical [28, 8] and pulse [4, 20, 35] movement of the trait optimum. This effect is due to allele frequency increases driven by displacement of the trait optimum, not balancing selection, and is highly sensitive to the assumed model of environmental change e.g. the amplitude and frequency of environmental cycles [8]. Thus, while it is already known that fluctuations can greatly increase *h*^2^, the issue is that special types of fluctuations are required to obtain this result [39]. If, as these studies suggest, we must assume rapid deterministic change in the trait optimum, then it is doubtful that fluctuations help to explain high heritability.

Here, instead of studying situational optimum changes (directional/cyclical/ pulse), we only allow undirected stochastic optimum fluctuations that accumulate slowly (compared to *V*_*e*_), and which are presumably commonplace in nature. Crucially, the optimum does not jump around trait space independently of its history (white noise), but rather follows a random walk, specifically an Ornstein-Uhlenbeck (OU) process. The change in trait optimum in any given generation is assumed to be small, but those changes (slowly) accumulate and eventually present an evolutionary challenge. OU trait optimum models are foundational in phylogenetic comparative methods [14], and have a similar interpretation to the model we develop here. That is, fitness depends on many different environmental factors many of which are continually changing. As a result, the optimal trait value wanders through trait space in unpredictable ways, albeit with constraints represented by the OU “restoring force”.

Apart from environmental fluctuations we follow classical Latter-Bulmer assumptions [30, 7], namely a single additive trait under Gaussian stabilizing selection, perfect linkage equilibrium and biallelic loci, although unlike LB we do account for genetic drift [11]. Thus, the model developed here can be thought of as LB with a random walk optimum.

Our model, (henceforth the “fluctuating optimum model”) behaves qualitatively differently to its stationary optimum counterpart even though the optimum only changes slowly. As the optimum changes, allele frequencies adjust to chase the optimum, and hence so does the additive genetic variance *V*_*g*_. But *V*_*g*_ also controls the rate at which the population chases the optimum. Thus, we have a feedback: *V*_*g*_ influences the time-course of selection, which in turn influences *V*_*g*_. Using a combination of simulations and mathematical analysis, we show that the heritability predictions resulting from this feedback are more consistent with observations and also less sensitive to evolutionary parameters than the classical LB predictions.

## 2 Methods

### 2.1 Stabilizing selection in a constant environment

We first briefly review the LB model [30, 7]. A quantitative trait with value denoted *z* is determined by additive contributions from *L* bi-allelic diploid loci, where individuals with trait value *z* have relative fitness

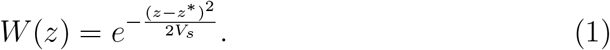

Here *z*^*^ is the trait optimum and the selection strength parameter *V*_*s*_ is assumed to be much larger than the total trait variance in the population.

Assuming linkage equilibrium between loci, the allele frequency change due to selection obeys the differential equation [2, 41]

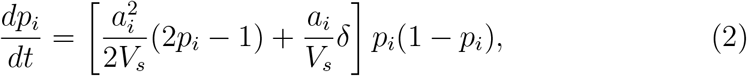

where *p*_*i*_ is the frequency of the focal allele at locus *i*, 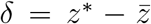 is the displacement of the trait mean 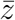 from the optimum, and *a*_*i*_ is the effect on *z* of switching one copy of a gene from the non-focal to the focal allele. Eq. (2) is conventionally derived assuming a normal trait distribution [2, 39]. In Appendix A we show that Eq. (2) also holds without normality. This is important because in some parameter regimes the number of segregating loci is too small to assume normality.

The LB model assumes the population is at the optimum (*δ* = 0). The remaining term in Eq. (2) is then negative for 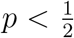 and positive for 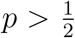 implying disruptive selection on allele frequency. This effect is sometimes also called “stabilizing selection” [2] but we will use “disruptive selection” to clearly distinguish it from the stabilizing selection model that also has a directional selection component when *δ*≠ 0.

The LB model thus has a selection coefficient

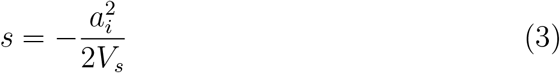

acting to purge low frequency alleles, including new mutations. This is counteracted by a mutation influx of *µ* per individual per locus per generation. In mutation-selection balance, minor alleles segregate at frequency *p*_*i*_ ≈ *µ/s*, and so the total additive genetic variance is given by [30, 7]

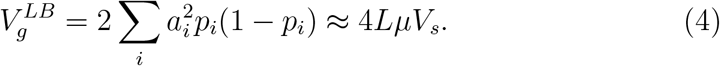

This is the same prediction as the house of cards model [36, 21].

Eq. (4) has the nice property of being independent of the effect sizes *a*_*i*_. In the model developed below, things are not so simple. To improve tractability we assume that effect size magnitudes are identical at all loci |*a*_*i*_| = *a*. We also assume forward and backward mutation rates are equal, which implies that the expected rate of change in *p*_*i*_ due to mutation is *µ*(1 − 2*p*_*i*_).

### 2.2 Random walk optimum fluctuations

We now relax the *δ* = 0 assumption and allow the optimum to move. Specifically, we assume 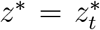 follows an Ornstein-Uhlenbeck (OU) process described by the stochastic differential equation (SDE)

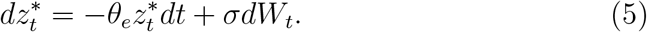

Here *θ*_*e*_ is a restoring force pulling 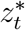 back to its long-run average, assumed without loss of generality to be *z*^*^ = 0, and *σ*^2^ is the variance in 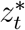 created each generation. If *θ*_*e*_ = 0, Eq. (5) is a Brownian motion (BM) process.

As the optimum moves through trait space, the magnitude of the optimum deviation 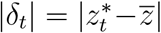 grows (on average), eventually becoming large enough that directional selection (second term in Eq. (2)) causes allele frequency shifts across all polymorphic loci. These shifts drive 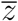 towards 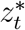 [15, 35]. The rate of pursuit depends on *V*_*g*_, and is given approximately by (Appendix B; [29, 4, 15])

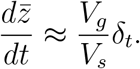

Consequently, *δ*_*t*_ approximately obeys the SDE

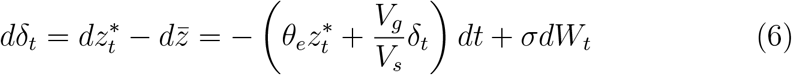

Eq. (6) is not an OU process for *δ*_*t*_, altough it approximately behaves like one for the purpose of determining steady-state *V*_*g*_. Eq. (6) differs from an OU process in two ways. First, the restoring force *V*_*g*_*/V*_*s*_ depends on time [4]. In the current model, the optimum deviations *δ*_*t*_ remain small enough that we can approximate *V*_*g*_ as time-independent (Appendix C). The time dependence of *V*_*g*_ is retained in our simulations (Sec. 2.3).

Second, the 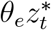 term implies Eq. (5) and Eq. (6) are coupled. Fortunately, the only property of the *δ*_*t*_ process we need is the temporal autocovariance (the covariance of *δ*_*t*_ with itself at other times). This we can compute from the two dimensional 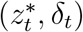 process defined by Eqs. (5)-(6) and the approximation of constant *V*_*g*_ as (Appendix B)

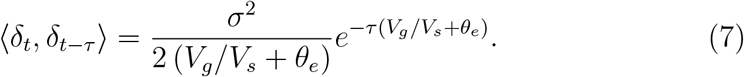

This is the same autocovariance function we would have if *δ*_*t*_ was an OU process with restoring force 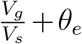 and mean *δ* = 0. Thus, *θ*_*e*_ effectively behaves like an additional restoring force on *δ*_*t*_. Eq. (7) says that the autocovariance decays exponentially with timescale 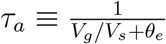.

### 2.3 Simulation

We simulate a discrete-time Wright-Fisher model using Eq. (2) to incorporate selection. Mutations are generated as a Poisson process at rate *µ* per locus per individual, with equal back/forward rates between the two possible alleles. When a mutation creates a new polymorphism at locus *i*, the new allele is assigned a trait effect *a*_*i*_ = ±*a* with with equal +*/*− probability. The behavior of 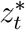 in Eq. (5) is approximated in discrete time as 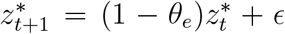 where *ϵ* is a normal random variable with mean 0 and variance *σ*^2^. To reach mutation-selection-drift balance, each replicate population is simulated for 10*N* generations, then *V*_*g*_ = 2*a*^2^∑ _*i*_ *p*_*i*_(1 − *p*_*i*_) is reported from the last generation.

### 2.4 Parameters

Our parameter choices are summarized in Table 1, with some explanation provided below. We choose our trait units such that *V*_*e*_ = 1, which means *V*_*g*_, *V*_*s*_, *a*^2^ and *σ*^2^ are all measured relative to *V*_*e*_. In particular, the narrowsense heritability becomes

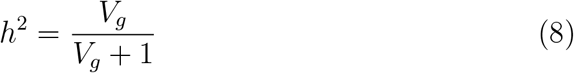

**Table 1:** Summary of parameters and their assumed values.

| Parameter | Symbol | Value(s) |
| --- | --- | --- |
| Environmental variance | $V_e$ | 1 |
| Selection strength | $V_s$ | 5, 20 |
| Mutation rate | $\mu$ | $6.6 \times 10^{-6}/\text{kb}$ |
| # of contributing loci | $L$ | $\leq 10^3$ |
| Effective population size | $N$ | $10^3, 10^4$ |
| Mutation effect size | $a^2$ | $\leq 10^{-1}$ |
| Fluctuation intensity | $\sigma^2$ | $\leq 10^{-2}$ |
| Env. restoring force | $\theta_e$ | $\leq 5 \times 10^{-3}$ |

In addition to the canonical *V*_*s*_ = 20 [36] we also include a stronger selection *V*_*s*_ = 5 scenario following [26, 21]. We assume an upper bound of *a*^2^ ≤ 0.1*V*_*e*_ on allele effect sizes [32].

In most of our simulations we assume *N* = 10^4^ for the drift-effective population size. This ensures *Ns >* 1 so that disruptive selection is not swamped by genetic drift. For instance, even relatively small effect sizes *a*^2^ = 0.01 and weaker stabilizing selection *V*_*s*_ = 20 would still have *Ns* = 2.5. We also include *N* = 10^3^ to explore the effects of stronger drift.

We use the mutation rate *µ* = 6.6×10^−6^/kb from a recent *D. Melanogaster* mutation accumulation experiment [18]. We assume each locus is a kilobase block, roughly corresponding to a gene, rather than modeling mutations at individual base pairs.

With the above parameter values, we need *L* = 10^3^ (= 10^6^bp) to reach 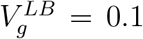 (the lower end of the empirical range) in the LB model under strong selection (*V*_*s*_ = 5). Since *L* ≥ 10^3^ implies an enormous genomic footprint for each trait [18], and we wish to demonstrate that it is not necessary to assume such large *L* values, we assume *L* ≤ 10^3^. The mutational heritability (influx of heritability due to mutation) is then at most 4*Lµa*^2^ = 2 × 10^−3^ (i.e. assuming *L* = 10^3^ and *a*^2^ = 0.1), on the higher end of empirical estimates but not implausibly large [21].

We assume slow environmental change *σ*^2^ ≤ 10^−2^. Thus, the fastest variance accumulation we allow is 1% of *V*_*e*_ per generation.

We constrain *θ*_*e*_ based on the implied trait optimum variance. Assuming *θ*_*e*_ > 0, the steady state variance implied by Eq. (5) is 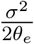 — we assume this variance is ≥ *V*_*e*_. In other words, the variability in trait optima the population experiences over evolutionary time (thousands of generations) is at least as big as the trait variability in one generation. Given our other parameter constraints this implies *θ*_*e*_ ≤ 5 × 10^−3^.

### 2.5 Data availability

Code and data used to generate figures can be accessed at github.com/jasonbertram/fluctuating optimum

## 3 Results

### 3.1 Simulation results: fluctuations greatly increase heritability

Random walk optimum fluctuations dramatically change the heritability predictions of simple quantitative genetic variation models. Fig. 1 shows an example of the dynamics involved. When the trait optimum is stationary (*σ*^2^ = 0), new mutations are deleterious, constraining them to low frequencies. With a fluctuating optimum (*σ*^2^ *>* 0), selection coefficients of new mutations are sometimes positive depending on *δ*_*t*_, releasing mutations to higher frequencies and even enabling fixation. As a result, *V*_*g*_ is substantially elevated in the presence of fluctuations. Fluctuations also induce much stronger variability in *V*_*g*_ over time, though *h*^2^ is almost never smaller than the *σ*^2^ = 0 case.

**Figure 1:**
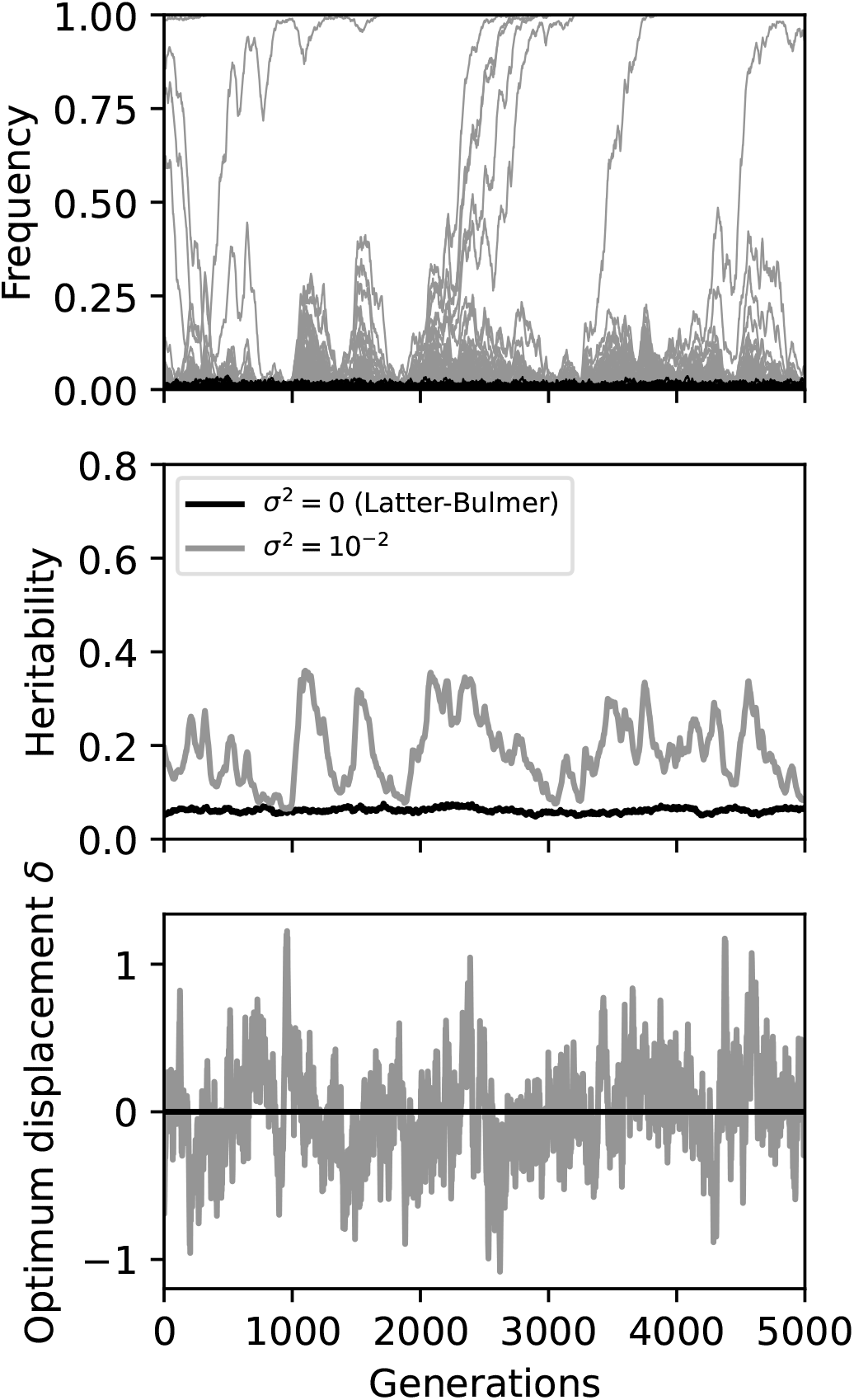
Example simulation of the fluctuating optimum model (*σ*^2^ = 10^−2^; grey) and constant optimum model (*σ*^2^ = 0; black) showing the strong increase of genetic variation in the presence of optimum fluctuations. Other parameters: *N* = 10^4^, *a*^2^ = 10^−1^, *θ*_*e*_ = 0, *V*_*s*_ = 5, *L* = 1000.

This behavior occurs without requiring rapid environmental change or particularly large accumulated optimum displacements. In Fig. 1, variance only accumulates at 1% of *V*_*e*_ per generation. The standard deviation of *δ*_*t*_ is ≈ 0.3, implying a typical directional selection coefficient of *a*|*δ*|*/V*_*s*_ ≈ 0.02. At most |*δ*_*t*_| occasionally reaches ∼ 1 giving *a*|*δ*|*/V*_*s*_ ≈ 0.06. These numbers assume large effect sizes (*a*^2^ = 0.1) and strong stabilizing selection *V*_*s*_ = 5. By comparison, under the same assumptions, the corresponding disruptive selection coefficient is *s* = *a*^2^*/*2*V*_*s*_ = 0.01, a number regarded as plausible based on empirical estimates of *V*_*s*_ [21].

Fig. 2 shows heritability predictions over many replicate populations as a function of *σ*^2^ assuming large effect sizes (*a*^2^ = 10^−1^), weak drift (*N* = 10^4^) and no environmental restoring force (*θ*_*e*_ = 0). Even for fluctuations as small as *σ*^2^ = 10^−3^ (0.1% of *V*_*e*_ per generation), the increase in the predicted average heritability is substantial. The effect gets bigger as *σ*^2^ increases, pushing *h*^2^ comfortably above 0.1 even for the challenging case of *V*_*s*_ = 5 and *L* = 100 where 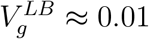. The increase in *h*^2^ is noticeably smaller for *σ*^2^ = 10^−4^, implying *σ*^2^ ∼ 10^−3^ is needed for optimum fluctuations to have a meaningful impact on *h*^2^ in our simulations.

**Figure 2:**
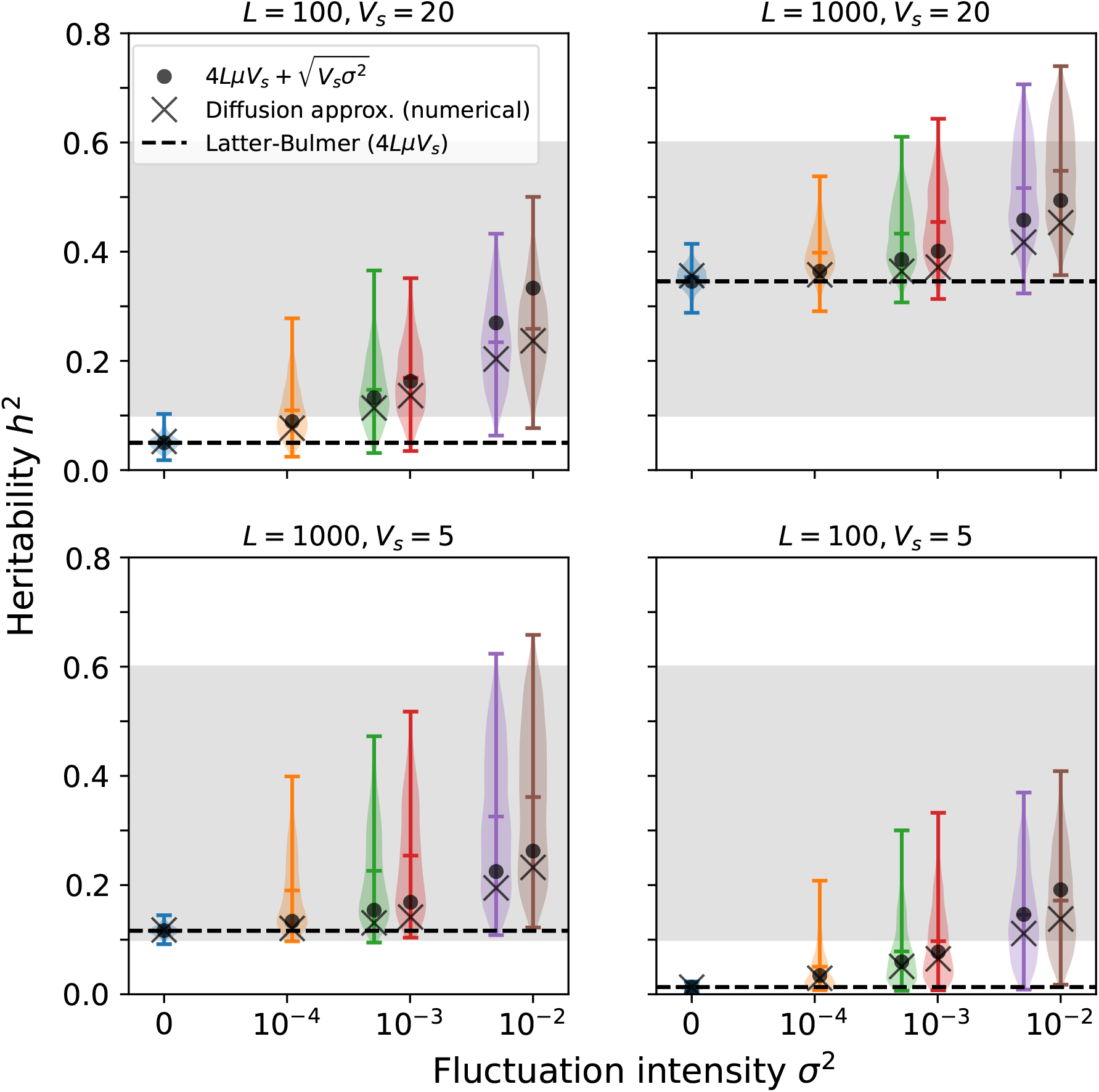
Simulated heritability in the random walk optimum model (violinplots of 10^3^ replicate populations) compared with the constant-optimum Latter-Bulmer prediction (dashed line). Region of typical heritability observed empricially 0.1 ≤ *h*^2^ ≤ 0.6 is shaded. Theoretical predictions are also shown (Sec. 3.3). Other parameters: *N* = 10^4^, *a*^2^ = 10^−1^, *θ*_*e*_ = 0.

Higher variance in heritability is also apparent in Fig. 2. As a result, *h*^2^ is frequently much higher than its mean, resulting in *h*^2^ values well above 0.1. This variance also implies *h*^2^ values frequently smaller than the mean, though it can be seen that values as low as the *σ*^2^ = 0 LB case are rare.

The results in Fig. 2 are robust to the environmental restoring force *θ*_*e*_. Under our parameter choices (Sec. 2.4) the pattern in Fig. 2 barely changes (Figs. S4-S5).

In order for fluctuating selection to influence frequency trajectories and have the effect shown in Figs. 1-2 we require strong selection on individual alleles relative to drift. Increasing the intensity of drift by setting *N* = 1000 weakens or eliminates the heritability increase caused by optimum fluctuations (Fig. S1). We discuss this strong selection requirement further below.

The effect size *a* also matters for our results, although the dependence of heritability on *a*^2^ is highly nonlinear. Reducing effect size to *a*^2^ = 10^−2^ (Fig. S3) has a big impact whereas reducing it to *a*^2^ = 0.4 has almost none (Fig. S2). To understand this counterintuitive behavior and the fluctuating optimum model’s heritability predictions more generally we now turn to model analysis.

### 3.2 Feedback between fluctuations and adaptation rate

Fluctuations in the present model are essentially continual changes in *δ*_*t*_, yet the model’s behavior depends crucially on the extent to which *δ*_*t*_ is temporally consistent. The key quantity is the temporal autocovariance Eq. (7).

To illustrate the importance of *δ*_*t*_ autocovariance, suppose we had none, replacing Eq. (6) with white noise: *δ*_*t*_ = *σdW*_*t*_. Now, assume *p* ≪ 1 (due to disruptive selection) and consider a time increment Δ*t* short enough that the *p*(1 − *p*) ≈ *p* factor in Eq. (2) is approximately constant. Then, from Eq. (6), the accumulated allele frequency change Δ*p* due to *δ*_*t*_ follows a Brownian motion process with variance (*ap/V*_*s*_)^2^*σ*^2^ per generation. In particular, after Δ*t* = 1*/s* generations, we have 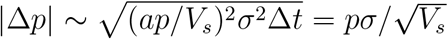. Thus we have |Δ*p*| ≪ *p* in our parameter regime because *σ*^2^*/V*_*s*_ ≤ 10^−2^*/V*_*s*_ ≤ 2 ×10^−3^. Over this same time Δ*t* = 1*/s*, disruptive selection induces a much larger change of order Δ*p* ∼ *p*. Thus, without autocovariance, fluctuations would barely perturb disruptive selection.

Unfortunately the effect of autocovariance in *δ*_*t*_ is hard to analyze because Eq. (7) depends on *V*_*g*_, which is what we want to predict. We cannot avoid this *V*_*g*_ dependence because it strongly affects the autocovariance. This is apparent from the ratio of autocorrelation timescale *τ*_*a*_ = *V*_*s*_*/V*_*g*_ (setting *θ*_*e*_ = 0 for illustration) to disruptive selection 1*/s* timescale

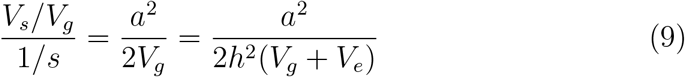

If *V*_*g*_ = *V*_*e*_ and *h*^2^ = 0.5 (high heritability), then the above ratio is ≪ 1 i.e. autocovariance decays rapidly. At the other extreme, if *V*_*g*_ = 10^−2^ and *h*^2^ = 10^−2^ (low heritability), then Eq. (9) evaluates to ≈ 5, implying little autocovariance decay over the disruptive selection timescale.

Thus we have a feedback. *V*_*g*_ controls the rate of adaptation, which influences *δ*_*t*_ and hence the fluctuations in selection coefficients, which in turn influences allele frequency dynamics and *V*_*g*_. This feedback is the main reason for the highly nonlinear parameter dependence of the fluctuating optimum model.

### 3.3 Diffusion approximation

We now summarize our approach for analyzing the steady state in the fluctuating optimum model and obtain a formula for *V*_*g*_. We proceed in two steps. First, we ignore the *V*_*g*_ feedback and compute the distribution of allele frequencies for fixed *V*_*g*_. Second, we use a self-consistency condition to resolve the feedback and solve for *V*_*g*_.

In the first step, we average our model over the autocovariance timescale *τ*_*a*_ = *V*_*s*_*/V*_*g*_. Since *δ*_*t*_ is approximately independent of 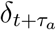, this averaging yields an uncorrelated diffusion approximation that captures the effects of autocovariance in its parameters (Appendix E; [33, 37]). This diffusion approximation is summarized by the mean rate of change in *p*_*i*_

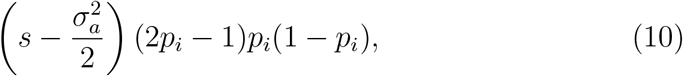

and the rate of variance accumulation

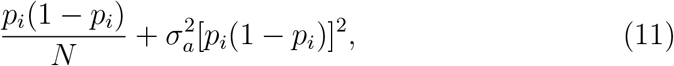

where

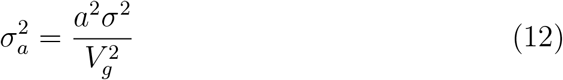

represents the influence of fluctuations in *δ*_*t*_ including its autocovariance.

Eqs. (10)-(11) are similar to single-locus fluctuating selection models [19, 33, 22] with some important differences. The 2*p* − 1 factor in Eq. (10) arises from stabilizing selection and is thus absent in single-locus models. In single-locus models 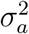 is a model parameter [33, 19]. Here 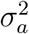 is not so simple because it depends on the unknown *V*_*g*_.

From Eq. (10)-(11), we calculate the distribution 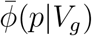 of allele frequencies for given *V*_*g*_ at mutation-selection balance (Appendix E)

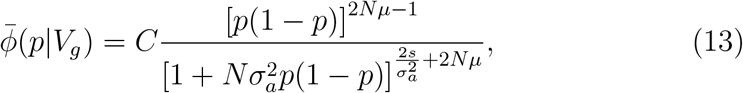

where 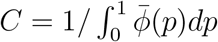 is a normalization constant.

In the limit *σ*^2^ → 0 we retrieve the stationary optimum distribution [11]

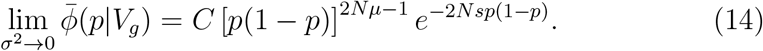

Compared with the neutral distribution *C*[*p*(1−*p*)]^2*Nµ*−1^, Eq. (14) introduces an exponential decay term that displaces frequencies away from intermediate values. When disruptive selection is strong enough to overcome drift as we assume here (*Ns* ≫ 1), then *V*_*g*_ will be much smaller than the neutral case.

With fluctuations 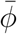 is also depressed at intermediate frequencies compared with the neutral distribution, but not to the same extent as Eq. (14). The factor 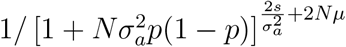 in Eq. (13) decays slower than exponential as *p*(1 − *p*) increases, approaching exponential only as fluctuations get vanishingly small and 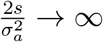.

This gives the first clue about the impact of optimum fluctuations. For 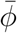 to decay slower than exponential, we need 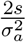 to be not much larger than 1. Thus, as an approximate heuristic we expect fluctuations to start to matter when

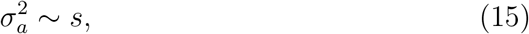

When this occurs, 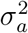 approximately cancels the disruptive selection coefficient *s* in the mean diffusion term Eq. (10).

To verify Eq. (15), we analyze allele frequency dynamics under the diffusion Eqs. (10)-(11) and confirm that 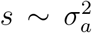 is required for fluctuations to start overpowering disruptive selection and release alleles to higher frequencies (Appendix G). For completeness, we also show that an analogous result holds in single-locus fluctuating selection models if there is purifying selection at strength *s* and the variance parameter is 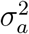.

At this point, however, the implications of Eq. (15) cannot be evaluated further because we do not know what *V*_*g*_ to use in 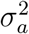 (Eq. (12)). For this we now proceed to our second step.

The second step is to find the steady-state *V*_*g*_ by applying a self-consistency condition to 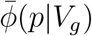. Assuming allele frequency dynamics are independent at each locus, 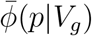 gives the steady-state distribution of allele frequencies among loci for a given *V*_*g*_. For consistency, the genetic variation implied by 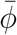 should therefore equal *V*_*g*_:

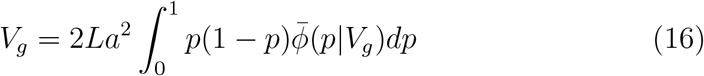

Thus, the steady-state *V*_*g*_ predicted by the diffusion model Eq. (10)-(11) is the solution of Eq. (16). This solution can be obtained numerically without difficulty since we have Eq. (13). We call this *V*_*g*_ the “diffusion approximation” for short in Fig. 2.

The diffusion approximation performs well when the number of available loci is small *L* = 100 (crosses in Fig. 2). However, when *L* = 1000 we underestimate *V*_*g*_. This is due the breakdown of the independence assumption in Eq. (16). As *L* increases, more alleles segregate at intermediate frequencies with allele frequency dynamics coupled through the optimum displacement *δ*_*t*_. This results in transient *V*_*g*_ spikes much larger than expected if loci were independent (Fig. 1). We do not attempt to address this limitation of Eq. (16) further here since we are most interested in the lower *L* scenarios where 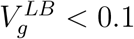. For larger *L* values our diffusion results can be interpreted as an underestimate.

In addition to solving Eq. (16) numerically, we derive an approximate formula for *V*_*g*_ (Appendix F) that is most accurate when fluctuations are just strong enough to start affecting genetic variation:

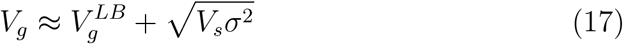

This approximation is also plotted in Figs.1-2 (large circles), and can be seen to do almost as well as the full numerical solution of the diffusion model (crosses).

Eq. (17) describes the state of the population at mutation-selection balance. When 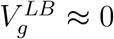, Eq. (17) is equivalent to 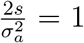 which we saw above is the requirement for fluctuations to matter (Eq. (15)). Therefore, in mutation selection balance, the population ends up approximately at this critical value.

Eq. (17) also allows us to estimate minimum *σ*^2^ required for the fluctuating optimum model to reach *h*^2^ = 0.1 in a case where the Latter-Bulmer model is failing (i.e. 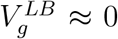). Namely *σ*^2^ ≈ 2 × 10^−3^, in agreement with our simulation-based estimate of needing approximately *σ*^2^ ∼ 10^−3^.

Eq. (17) starts to break down if selection on individual alleles becomes too weak or the mutational influx becomes very small (formally we require, 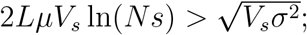 Appendix G), but above this point the contribution due to fluctuations 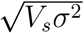 is approximately independent of *a* and *L*. This remarkable situation explains the extreme nonlinear dependence on *a* and the greatly reduced sensitivity to *L* in simulations.

### 3.4 Required mutational target sizes

We now address the standard critique of the Latter-Bulmer model (and house of cards approximation), and by extension our broader understanding of quantitative genetic variation [36, 21, 18, 39].

LB model predictions are often analyzed in terms of the *L* that would be needed to reach observed heritabilities. The reason for this is that LB model predictions depend only on *µ, V*_*s*_ and *L*, of which *L* is the least certain. *µ* can be measured fairly accurately by mutation accumulation [18]. *V*_*s*_ is less certain but we operate under the standard assumption that *V*_*s*_ *>* 20 is unlikely for many traits of interest [21].

Thus, in Fig. 3 we show the LB and fluctuating optimum model predictions as functions of *L*. Since our *µ* is taken from *D. Melanogaster* we express *L* as a proportion of the *D. Melanogaster* euchromatic genome (128Mb) following [18].

**Figure 3:**
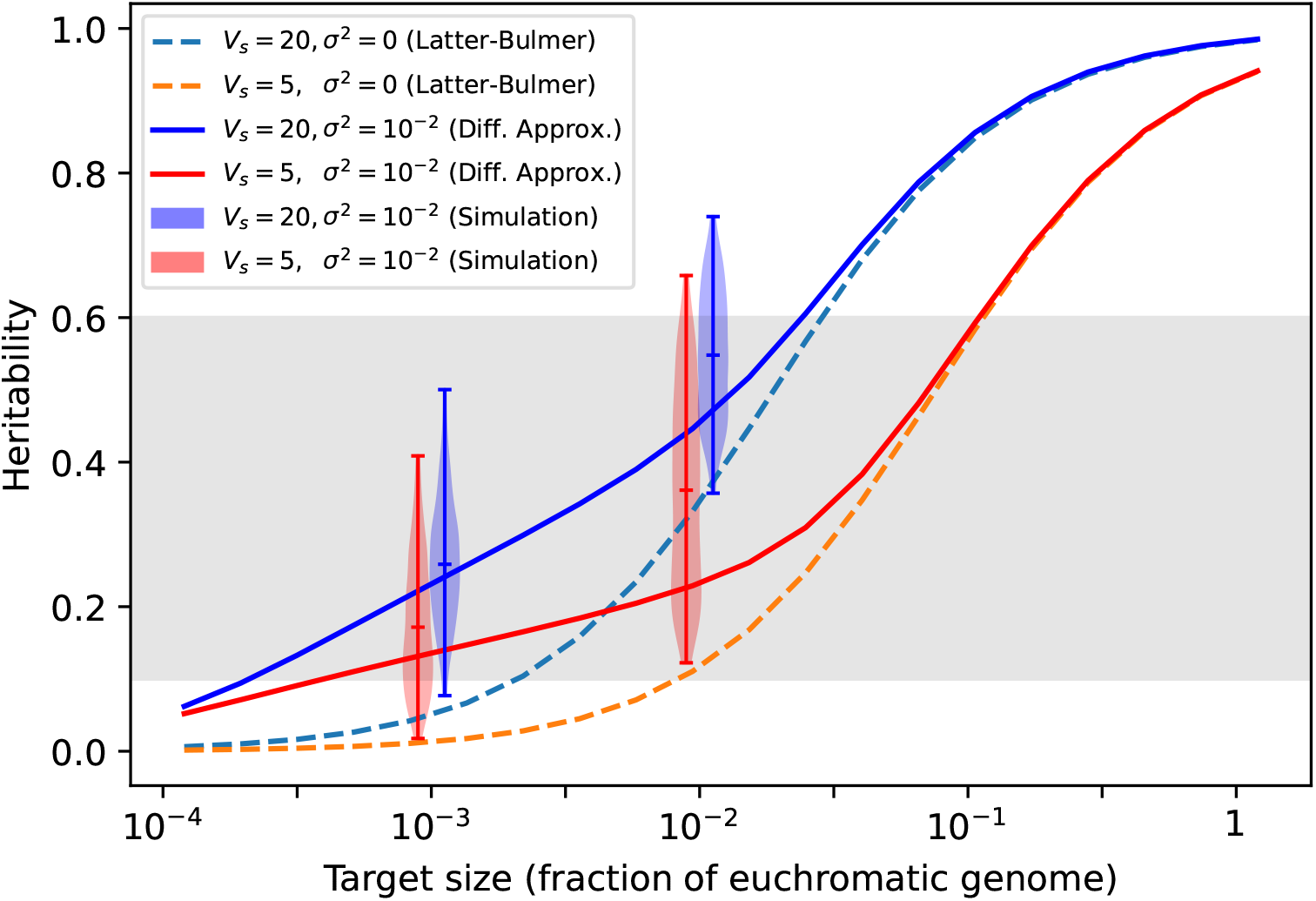
The mutational target size needed to reach *h*^2^ = 0.1 is reduced by more than an order of magnitude in the fluctuating optimum model (simulations in violinplot, diffusion approximation in solid lines) compared to the Latter-Bulmer model (dashed lines). Other parameters: *N* = 10^4^, *a*^2^ = 10^−1^, *θ*_*e*_ = 0.

Fig. 3 shows that the *L* needed to reach *h*^2^ = 0.1 in the fluctuating optimum model is reduced by more than an order of magnitude down to around 0.01% − 0.1% of total euchromatin. Note that for higher *L* values the fluctuating optimum and LB model predictions converge. This is again due to the feedback driving the change in heritability — if *h*^2^ is inherently large due to high mutational influx, the population is able to rapidly track the moving optimum and *δ*_*t*_ never grows large enough to change the structure of genetic variation. Thus the fluctuating optimum model is also robust to high *L* values, only contributing extra *V*_*g*_ where it is “needed”.

## 4 Discussion

The fluctuating optimum model developed here makes surprisingly different predictions from the static optimum LB model given that the only change is slow, undirected movement of the trait optimum. The LB model epitomizes basic intuitions about mutation-selection balance: stabilizing selection removes variation (the *V*_*s*_ factor in Eq. (4)) while mutation replenishes it (the *Lµ* factor in Eq. (4)). In this picture, strong stabilizing selection is a challenge for MSB models [21, 39], one that might be overcome by massive mutational target sizes [32].

In the fluctuating optimum model on the other hand, there is an additional component of *V*_*g*_ given approximately by 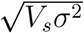, which makes up the bulk of *V*_*g*_ in the cases where the Latter-Bulmer heritability is close to zero. Remarkably, this expression is independent of both *L* and *µ* and hence the mutational input of new variation. The square-root dependence on *σ*^2^ implies that even small fluctuations have a big impact. Moreover, in contrast to LB where stronger selection strictly reduces *h*^2^, the fluctuating optimum model actually *requires* strong selection (relative to drift) to meaningfully increase *V*_*g*_.

From this perspective the LB model is a *σ*^2^ = 0 edge case unrepresentative of *σ*^2^ *>* 0 models even very close to it. Its predictions should therefore be treated with skepticism if there is reason to doubt that the optimum is static, which there typically will be.

It may seem obvious from Eq. (2) that even slow optimum movement can produce qualitatively different predictions. The optimum displacement |*δ*| only needs to be of order *a* (the trait contribution of a single allele) for the directional term 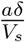 to be as important as the disruptive term 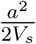. However, this reasoning is deceptive because it implicitly assumes that, like disruptive selection, the optimum displacement is consistent over time. This would require narrow assumptions about the environment that we certainly do not make in the fluctuating optimum model. Instead we assume stochastic optimum displacements that are presumably commonplace in nature. The cumulative effect of stochastic displacements is harder to evaluate. A fairly length analysis was needed (Sec. 3.3) to show that indeed the cumulative effect of random walk fluctuations can overcome disruptive selection.

Although the fluctuating optimum model’s predictions are substantially better than LB, it is also not sufficient to explain high heritabilities in nature. Apart from incorporating more realistic environmental assumptions, the fluctuating optimum model remains heavily simplified in other respects. A particularly big limitation is the absence of multiple traits and pleiotropy. The large mutation targets found in GWAS studies are not possible without pervasive pleiotropy [32]. Moreover, if every trait that appears to be under selection was independently and directly experiencing that selection it would imply unfeasibly strong selection on fitness [3]. Attempts to account for pleiotropy in simple heritability models have faced similar issues as the LB model. For instance, “apparent” selection models, which assume that stabilizing selection on individual traits results solely from pleiotropy between the trait and fitness, are unable to reproduce the observed intensity of selection and have pathological dependence on *N* [21]. Our model does nothing to address the essential question of pleiotropy.

Moreover, the parameter *θ*_*e*_ could in principle be larger than we allow, which would dampen the effect of fluctuations considerably. This would imply that the environment is so tightly constrained that the long-run variance in the trait optimum is smaller than the environmental trait variance *V*_*e*_, which would seem to represent an extremely tightly constrained environment. Regardless, the magnitudes of *θ*_*e*_ and also *σ*^2^ used here would ideally be supported emprically. Similar quantities are routinely estimated in the phylogenetic comparative literature but it is not apparent that existing estimates are applicable since these analyses span longer timescales than we have in mind and also rely on comparing the evolution of different species [14].

What we claim our results *do* show is that simple models of quantitative genetic variation can attain observed values of *h*^2^ without having to assume extreme mutational target sizes, contrary to long-standing consensus [21, 39, 18]. This finding is important because, if MSB is responsible for high trait heritabilities, we expect that even a simplified accounting of the dominant effects of mutation and selection would predict heritabilities that are at least the correct order of magnitude. Failing this we are faced with two options.

First, we could embrace the main alternative hypothesis to MSB: balancing selection (selection that preserves polymorphism). Currently there is no empirical support for balancing selection being sufficiently widespread to explain consistently high heritabilities [32]. Moreover, from a theoretical perspective, models of balancing selection almost always require large selection coefficients and organism-specific assumptions about the form of selection (e.g. particular life histories), raising the question of how balancing selection could explain consistently high heritability even in principle [6]. On the other hand, some manifestation of mutation-selection-balance is almost certainly present to a substantial extent in every trait/species.

This leaves the second option: the failure of simple quantitative genetic models indicates fundamental holes in our understanding of quantitative trait evolution [21, 39, 18]. Our findings suggest that perhaps one big hole is environmental change, which has either been omitted or modeled in ways that have not captured the potential generality and magnitude of its impact. Many aspects of our model could be made more realistic including allowing multiple traits and pleiotropy, linkage disequilibrium, and different effect sizes. We hope these will be pursued in future work. However, we emphasize that the mechanism driving increased heritability in the fluctuating optimum model is not likely to be sensitive to these factors. The essence of the variance-increasing effect captured by the fluctuating optimum model is a feedback between adaptation rate and selection coefficient fluctuations. We anticipate similar effects on heritability will occur in more complicated models provided the optimum moves in a comparable way i.e. a random walk with long-term variance greater than *V*_*e*_.

## Acknowledgements

ZS was funded by NSERC 2024-05237. This research was enabled in part by support provided by Compute Ontario (computeontario.ca) and the Digital Research Alliance of Canada (alliancecan.ca).

## Author contributions

JB: conceptualization, formal analysis and writing the manuscript. ZS: formal analysis in Appendix G.

## Appendices

## A Selection equation

The conventional derivation of Eq. (2) relies on assuming a normal distribution for *z* [2, 39]. Here we give an alternative derivation showing normality is not necessary provided *V*_*s*_ is large relative to the phenotpyic variance.

The selection equation in continuous time at locus *i* is

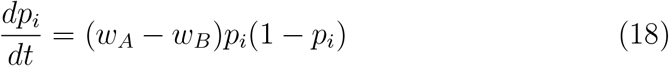

where *w*_*A*_ is fitness of the focal *A* allele, defined as an average

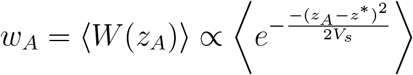

over all individuals with an *A* allele at locus *i* (similarly for *B*). We have

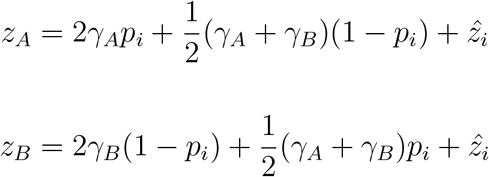

where 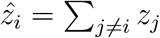. Moreover,

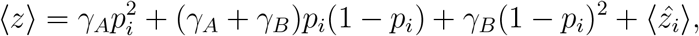

hence

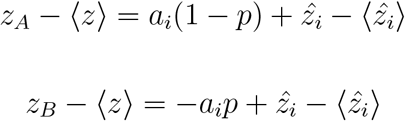

where 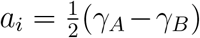. Since we assume linkage equilibrium, the distribution of genetic backgrounds for *A* and *B* are identical such that 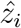 can be viewed as a random variable that is identically and independently distributed for the *A* and *B* alleles at the focal locus.

Assuming *V*_*s*_ is large compared to the phenotypic variance, the fitness function is well approximated by the leading term in its Taylor expansion, giving

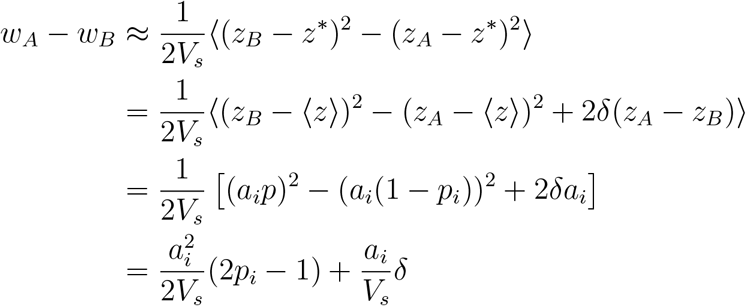

where 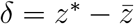. This is the required result.

## B Ornstein-Uhlenbeck process for *δ*_*t*_

Focusing on the directional pull to the optimum and ignoring the effects of drift and disruptive selection (which are negligibly weak by comparison), we have 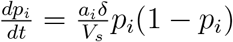 and so

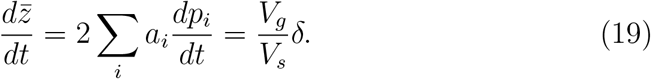

If there is an optimum restoring force such that *θ*_*e*_ *>* 0, we have the coupled system of stochastic differential equations

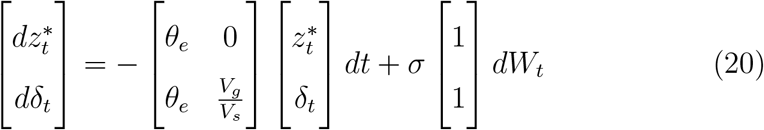

which, to leading order in the restoring force terms (i.e. ignoring 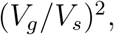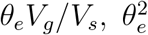 and higher order terms) has steady-state temporal covariance given by [13, Sec. 4.5.6b,d]

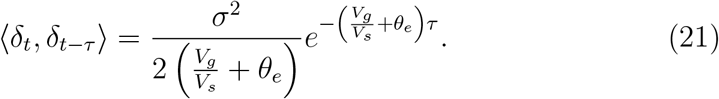

## C Approximate constancy of *V*_*g*_

The rate at which *V*_*g*_ changes is

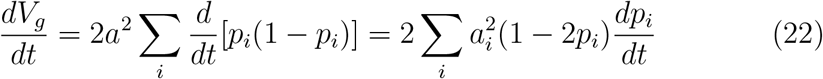

where

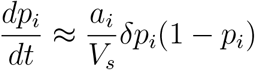

because allele frequency change is dominated by the directional component during a return to the optimum.

On the other hand, the rate at which the mean trait value changes is

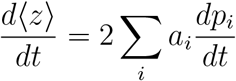

Comparing these rates, we see in most cases 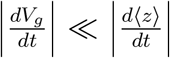. These sums differ only due to the presence of an additional factor of *a*_*i*_(1−2*p*_*i*_) in Eq. (22). By assumption 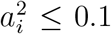. We also have |1 − 2*p*_*i*_| ≤ 1 and moreover, alleles at intermediate frequencies which make the largest individual contributions to 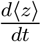 induce neglible change in *V*_*g*_. Additionally, all terms in 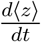 have the same sign, whereas the terms in 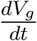 will often have opposite signs, partly cancelling each other out.

Therefore, over the autocorrelation timescale *τ*_*a*_, which is also the minimal timescale required for appreciable change in ⟨*z*⟩, the change in *V*_*g*_ will not be large enough to warrant the complication of treating its full time-dependence.

## D Averaging the autocovariance

Incorporating genetic drift into Eq. (2), we obtain

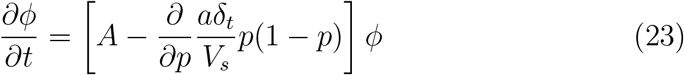

where we have introduced the operator

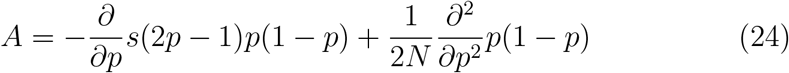

and *ϕ* = *ϕ*(*p, t*|*p*(0)) is the probability density function for an allele to have frequency *p* at time *t* given it had frequency *p*(0) at *t* = 0.

The stochastic variable *δ*_*t*_ obeys Eq. (6) and has autocovariance timescale *τ*_*a*_. We now remove autocovariance from Eq. (23) using the cumulant expansion [37], which effectively integrates the behavior of *δ*_*t*_ over the interval *τ*_*a*_ and gives an approximate “effective” uncorrelated diffusion model. This approximation requires two conditions:

1. Fluctuations effectively behave like uncorrelated pulses when viewed in time increments of *τ*_*a*_. Formally, we require [37, pp. 400]

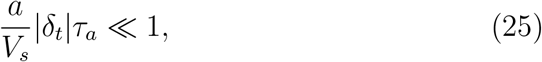

where

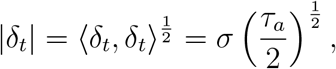

and so

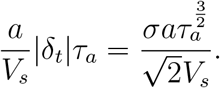

Now choose *V*_*s*_ = 5, *θ*_*e*_ = 0, *σ*^2^ = 10^−2^ and *a*^2^ = 0.1 i.e. the worst case choices in each parameter’s assumed range for satisfying Eq. (25). This gives

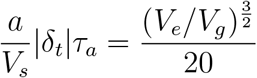

Thus, Eq. (25) is satisfied if 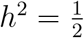, but starts to break down as *h*^2^ falls to 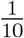. This is the worst case; for other parameter choices even lower values of *h*^2^ will satisfy Eq. (25).
2. Allele frequency change due to *A* (disruptive selection + drift) must be negligible over the interval *τ*_*a*_ [37, pp. 401]. This condition is not satisfied near *p* = 0, 1 where drift dominates over any selection, including directional selection due to optimum deviations. However, in practice this boundary behavior is not important because both the exact and approximated optimum fluctuations are negligibly small near *p* = 0, 1. What does matter is the component of *A* due to disruptive selection, which starts to exceed the change due to drift at the frequency scale 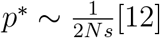. Formally, we require

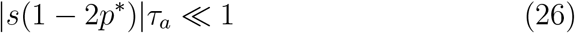

Similar to the first condition above, we assume without loss of generality that *θ*_*e*_ = 0 and *a*^2^ = 0.1, *V*_*s*_ = 5 and *N* = 1000. Then *p*^*^ ≪ 1 and Eq.(26) reduces to the timescale ratio in Eq. (9). Similar to Eq. (25), this condition is comfortably satisfied at *h*^2^ = 0.5, but starts to break down as we approach *h*^2^ = 0.1.

When the above conditions break down, the approximate averaged diffusion model underestimates the influence of optimum fluctuations since it effectively discards autocovariances that persist between the averaging intervals of length *τ*_*a*_. Treating fluctuations as though they truncate rapidly in the low *h*^2^ situation where they do not underestimates the *V*_*g*_ increase that they cause — some positive autocovariance is effectively being ommitted. But, in practice, since *h*^2^ is already close to zero, underestimating in this case *V*_*g*_ makes little difference.

Performing the cumulant expansion and only retaining the first fluctuation term [37], we obtain the diffusion equation

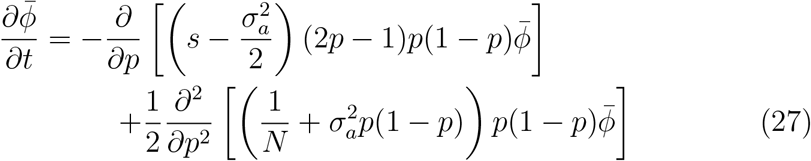

where 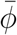 is *ϕ* averaged over the autocorrelation timescale and

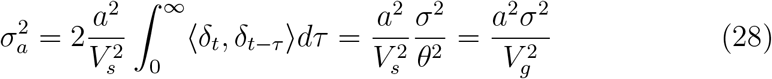

## E Steady state density

To derive the steady state density we first incorporate mutations into the above diffusion equation by adding *µ*(1 − 2*p*) to the mean term. The steady state solution of Eq. (27) is then obtained by setting 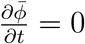 and integrating, which gives

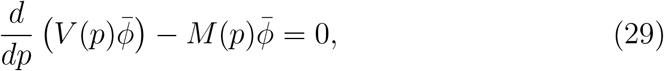

where

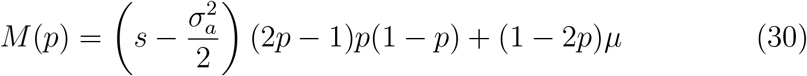

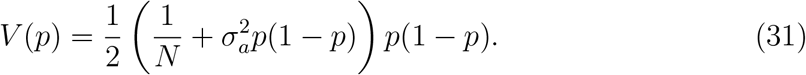

Note that the integration constant is zero (i.e. zero probability flux) due to the symmetry of the model about 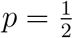. Multiplying Eq. (29) by 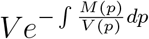 and integrating gives

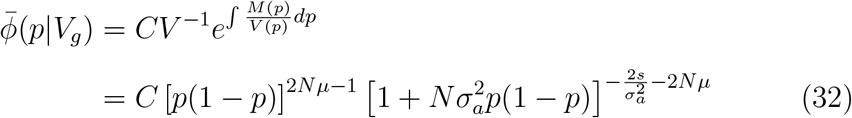

where *C* is a constant that ensures 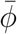 is a probability density i.e. 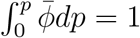.

## F Steady-state genetic variance

The distribution 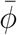 allows us to compute the steady-state additive genetic variance from Eq. (16). We were not able to obtain an exact expression for this integral in terms of elementary functions. Here we derive an approximation that works well in the parameter regime of interest.

Under the present parameters we have *Nµ* ≪ 1, and so 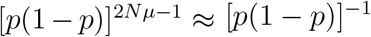. Hence the integrand is approximately

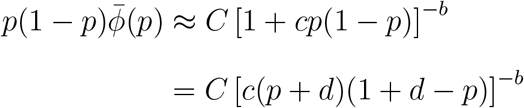

where 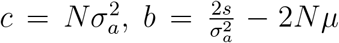 and 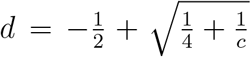. This function is symmetric about 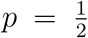, and since *b >* 0, the points *p* = −*d <* 0 and *p* = 1 + *d >* 1 are singular. 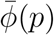 thus changes most rapidly near *p* = 0, 1. In particular, the factor (*p*+*d*)^−*b*^ changes much faster than the factor (1+*d*−*p*)^−*b*^ in the neighbourhood of *p* = 0. We approximate the integrand by introducing an effective exponent 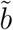 that satisfies

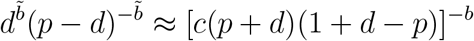

near *p* = 0. Matching the gradients of these functions gives

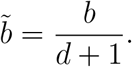

We now integrate:

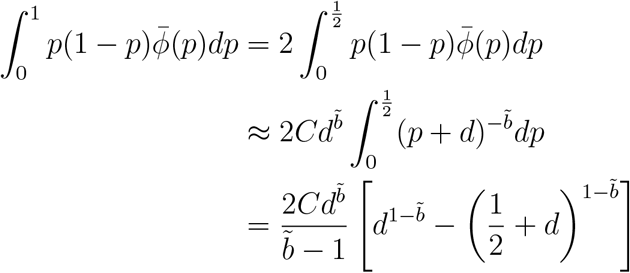

To evaluate the normalization constant *C* we use the fact that 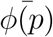 is singular at *p* = 0, 1. Compared with this divergence to ∞ the change in the 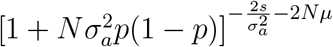 factor in 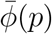 is small. Thus we approximately evaluate *C* as

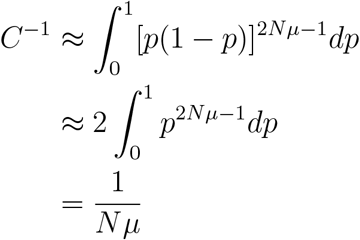

Therefore

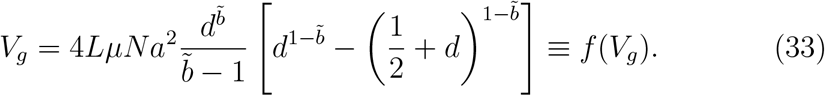

where *f* (*V*_*g*_) is a decreasing concave-up function. The equation *V*_*g*_ = *f* (*V*_*g*_) involves exponents of the form 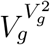 which makes analytical solution difficult.

Let 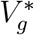 denote the solution of Eq. (33) i.e. the intersection of *f* (*V*_*g*_) with the one-one line. Evaluating the extremes of *f* (*V*_*g*_) assuming 0 *< σ*^2^ ≪ 1, we have

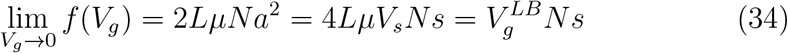

and

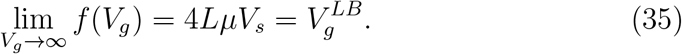

Hence

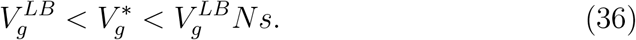

The upper bound in Eq. (36) is much greater than 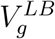 because *Ns* ≫ 1. Therefore, depending on the shape of *f* (*V*_*g*_), 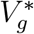 can be much greater than 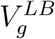.

We now derive an approximate expression for 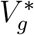.We first evaluate *f* (*V*_*g*_) at a trial value 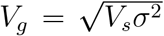 (this choice will be explained below), and we also assume 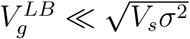. We have

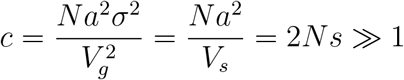

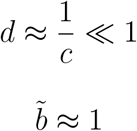

and so

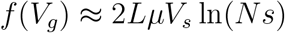

It follows that

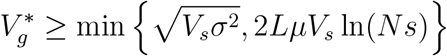

Assuming selection is sufficiently strong relative to drift ln(*Ns*) ≫ 1, we therefore have

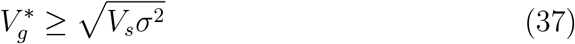

The trial value 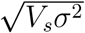 corresponds to *b* = 1, and *b* increases with *V*_*g*_ with greater *b* implying weaker influence from fluctuations (Sec. 3.3). Therefore, since 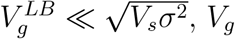 cannot be appreciably larger than 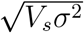, since any extra variation over and above 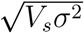 would also have to be generated by fluctuations. Consquently, 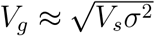.

Finally, we relax the assumption of small 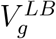 by assuming 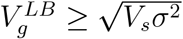. In this case, we consider a rescaled version of 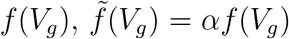, where *α* ≪ 1. Thus, 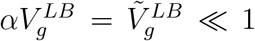. Intuitively this can be thought of as evaluating a different population with a lower value for *L*. Then

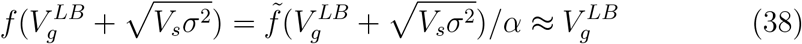

since 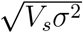 is the approximate fixed point of 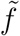 and 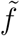 decays rapidly with increasing *V*_*g*_. It can also be seen geometrically that 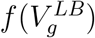 lies well above the fixed point of *f* (*V*_*g*_). Consequently, the fixed point must be close to 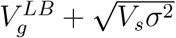.

## G Balance of forces on an allele frequency trajectory under fluctuating selection

Here we analyze the influence of disruptive and fluctuating selection on allele frequency dynamics. We begin with the single-locus model by Karlin and Levikson [22] before addressing the fluctuating optimum model.

In the Karlin-Levikson model, two alleles have fitness 1+*α* and 1+*β* where *α* and *β* are independent random variables drawn from two distributions with the same shape and variance *σ*^2^. Therefore,

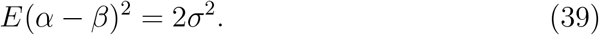

We slightly modify the model of [22], which assumes that *α* and *β* are identically distributed and assume that new mutations are expected to be deleterious on average. Mathematically,

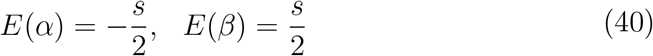

where *s* is analogous to the disruptive selection coefficient in the main text.

Huerta-Sanchez et al. [19] derived the diffusion limit for this model, which reduces to

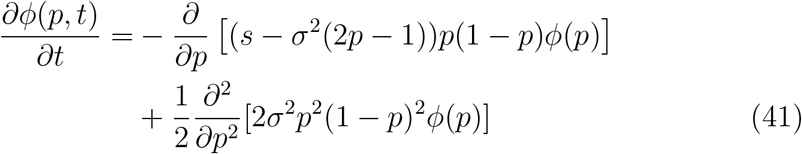

The SDE corresponding to Eq. (41) is

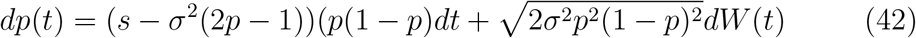

Integrating over a time interval Δ*t* that is short enough to prevent big changes in *p, p* is approximately constant with respect to *t* and we have a total allele frequency change of

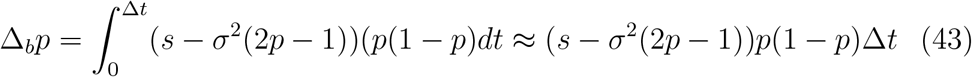

Therefore, the relative change in frequency due to the negative shift is:

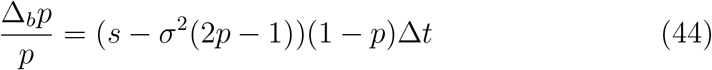

Denoting the relative change in frequency by *η*,

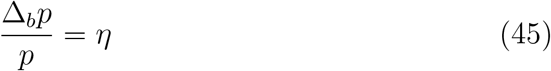

and solving for Δ*t*, we have:

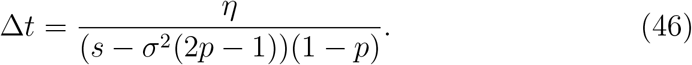

Next, we assess how much allele frequency change is caused by selective fluctuation during this time period. Given that we need to consider the stochastic component of Eq. (42), we use the Itô isometry property:

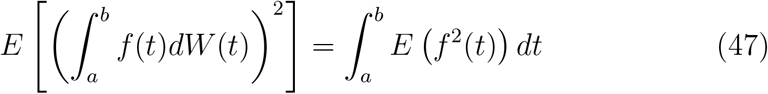

where *f* (*t*) is a random function. Applying Eq. (47) to the stochastic component of Eq. (42), we obtain

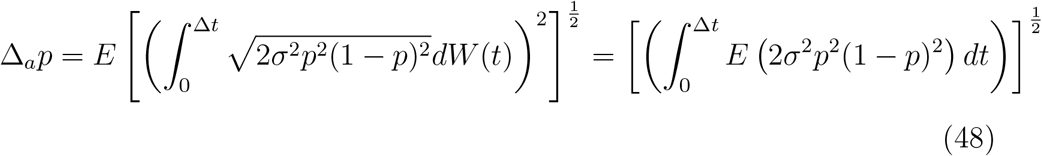

Assuming again that Δ*t* is small enough that *p* can be approximated as constant over time, we obtain

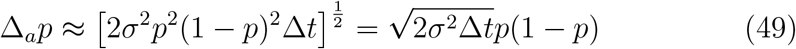

and hence

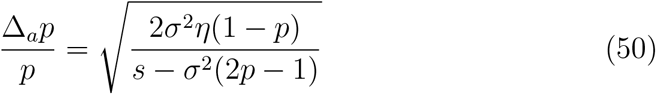

Focusing on low frequencies *p* ≪ 1 and setting *η* = 1 so that the Δ*p* is of order *p*, and further assuming 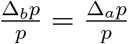, we finally obtain

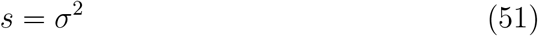

This result tells us what is needed for fluctuations to cause comparable allele frequency changes as the constant negative selection ocurring at strength *s*. When *σ*^2^ *< s*, negative selection has more influence, and when *σ*^2^ *> s*, selective fluctuation has more influence. Remarkably, this condition is frequency-independent.

We now apply the same procedure to the the diffusion model in the main text, given by Eq. (27). The corresponding SDE is now:

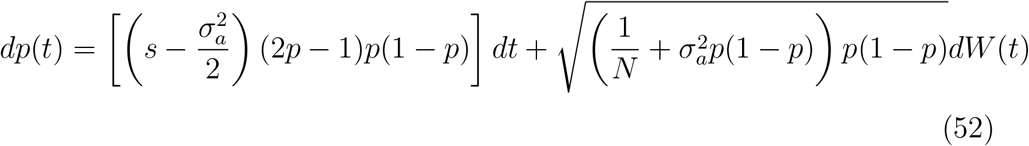

where 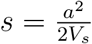 now denotes the disruptive selection coefficient.

The change in frequency due to directional part of the equation is similarly

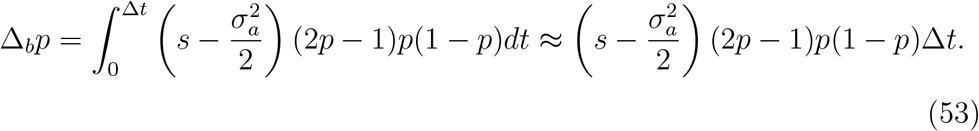

Assuming 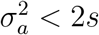, we have

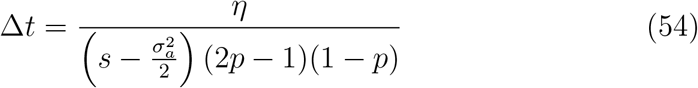

where *η* = Δ_*b*_*p/p*.

Next, we apply the Itô isometry property to the flucutating selection component:

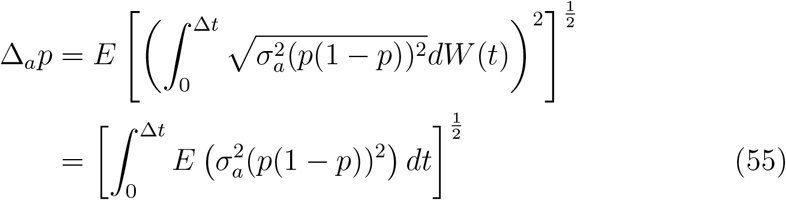

which similarly leads to

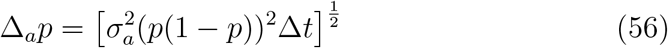

Again setting |Δ_*a*_*p/p*| = |Δ_*b*_*p/p*| and *η* = 1 we obtain the condition under which optimum fluctuations and disruptive selection have comparable influence, which is now frequency-dependent (i.e. we solve for the frequency at which this occurs)

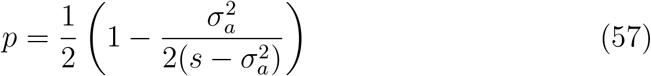

Thus, if 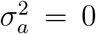, then 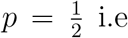 the frequency at which the mean diffusion term vanishes. As 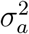 increases, *p* decreases and fluctuating selection is stronger over a growing interval of intermediate frequencies. At 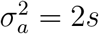, stabilizing selection vanishes again, this time over the entire frequency domain.

## Supplementary figures

**Figure S1:**
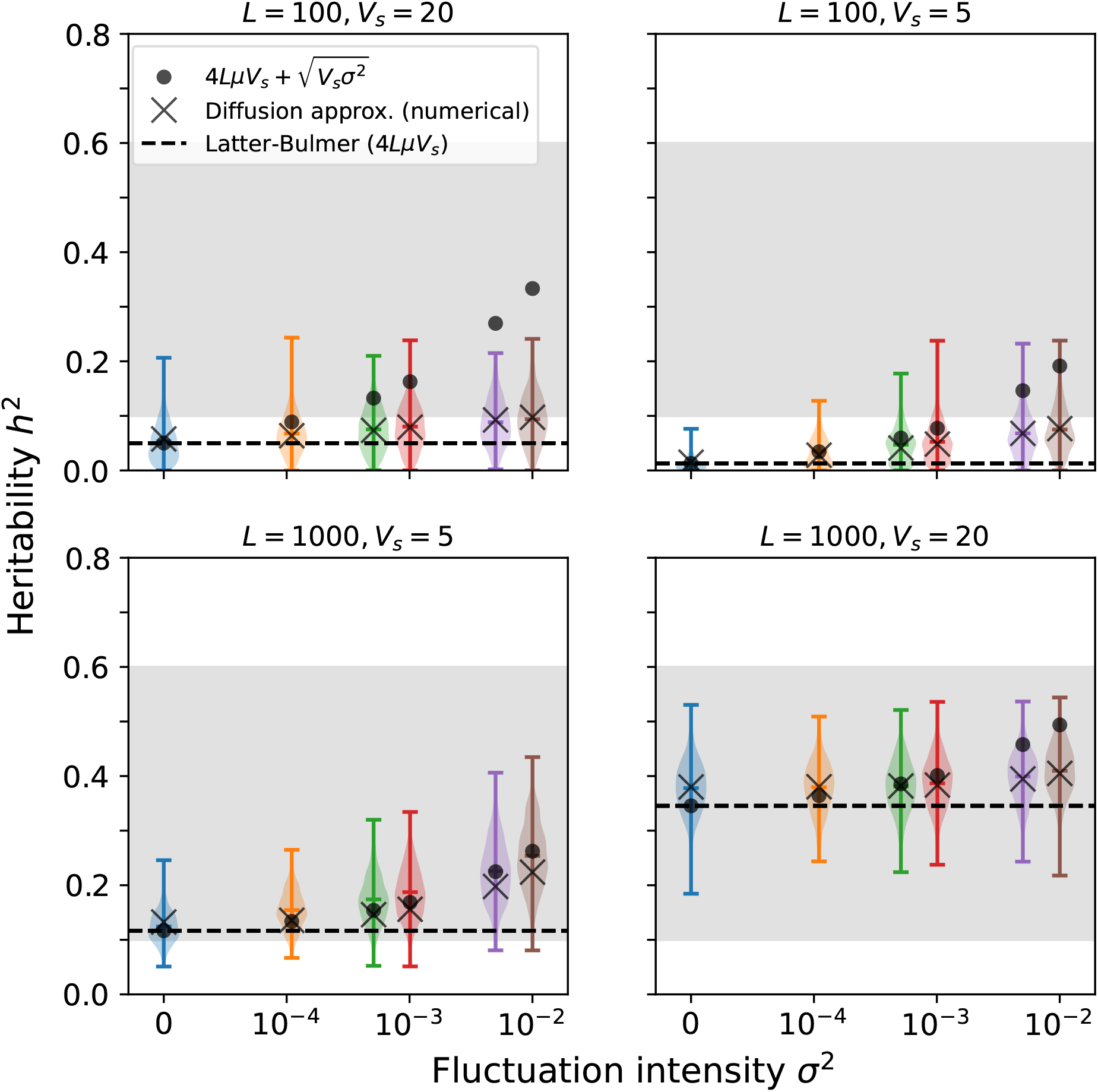
Same as Fig. 2 but with stronger drift *N* = 1000.

**Figure S2:**
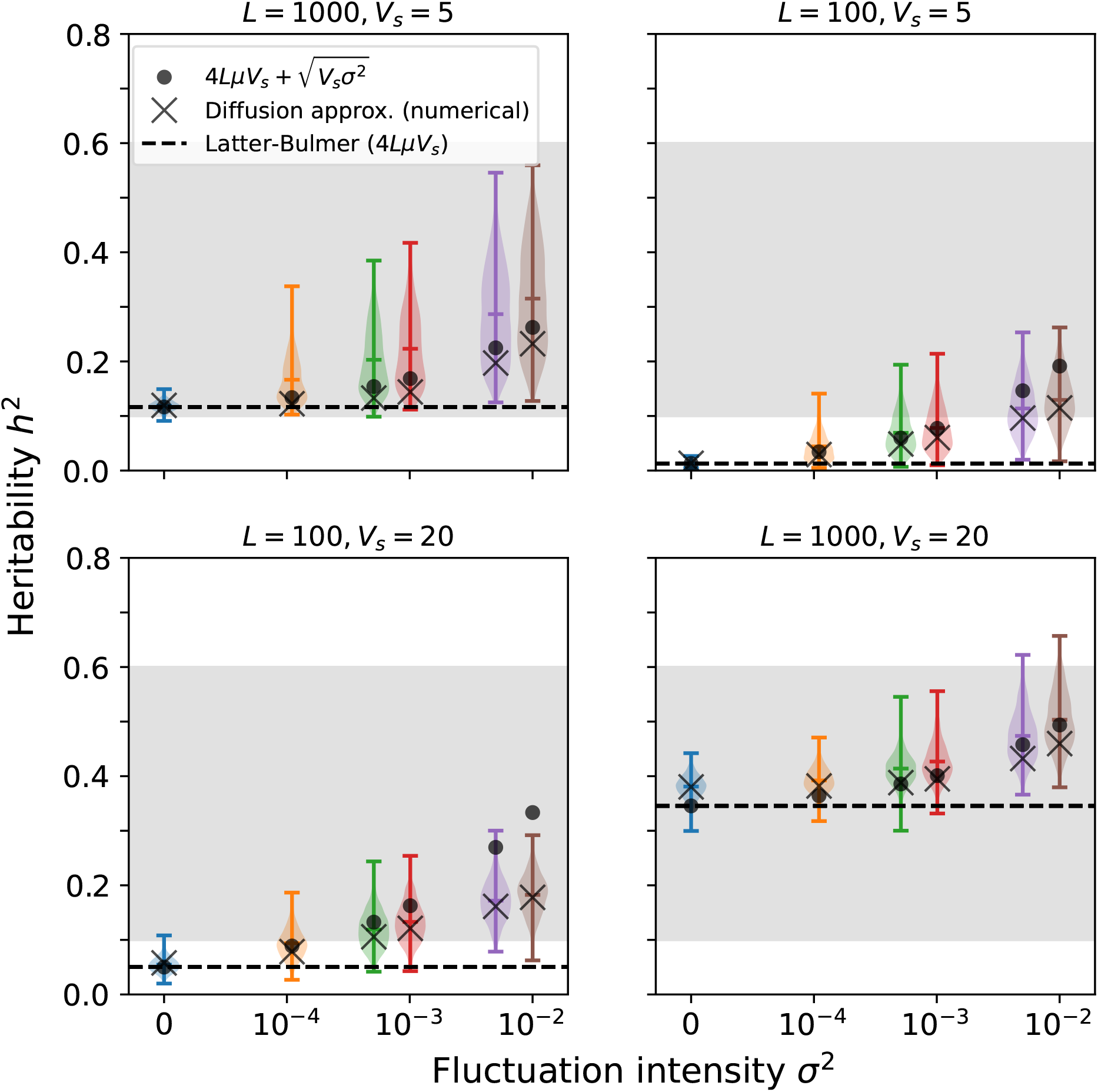
Same as Fig. 2 but with weaker allele effect size *a*^2^ = 0.04.

**Figure S3:**
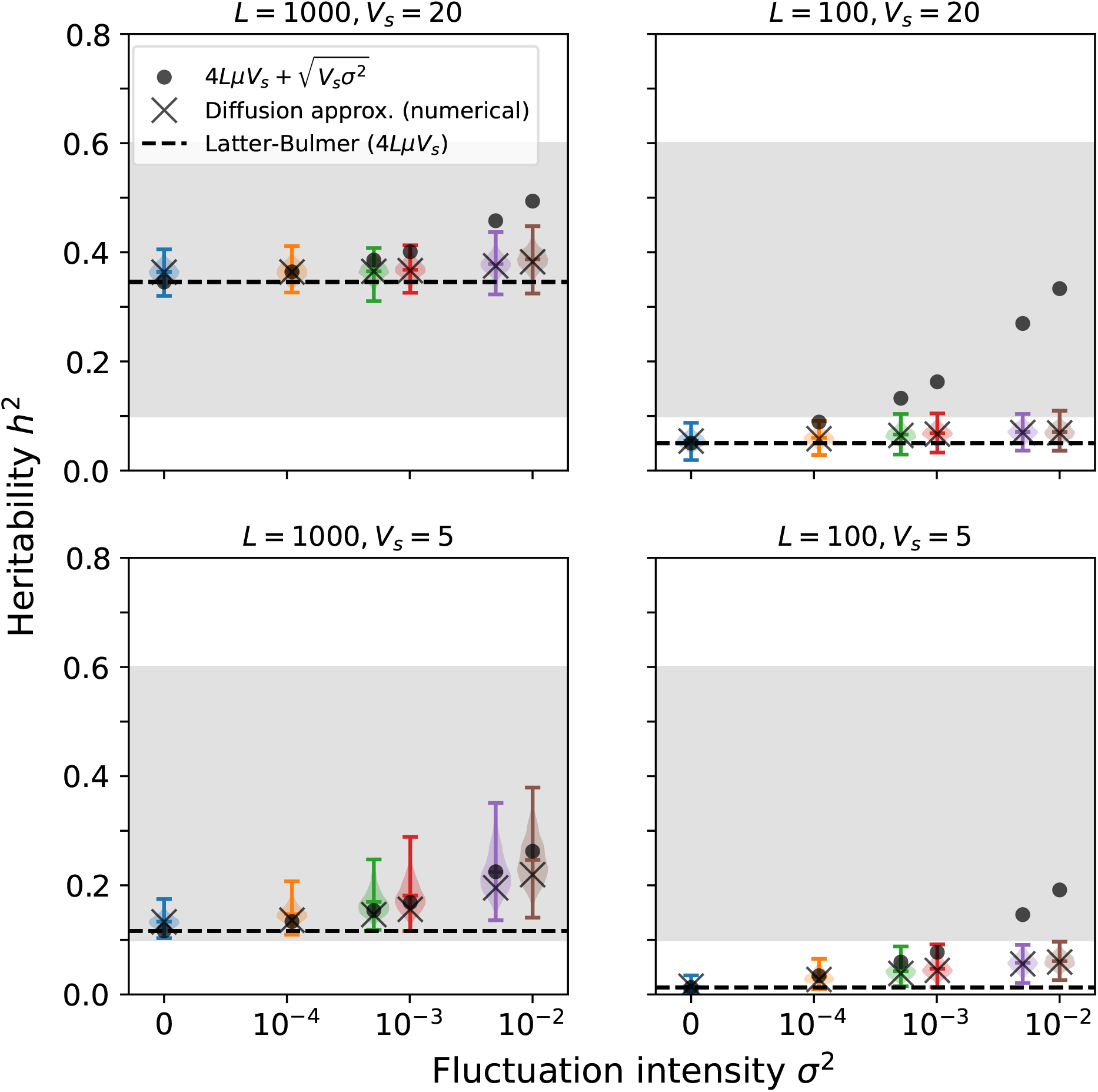
Same as Fig. 2 but with weaker allele effect size *a*^2^ = 0.01.

**Figure S4:**
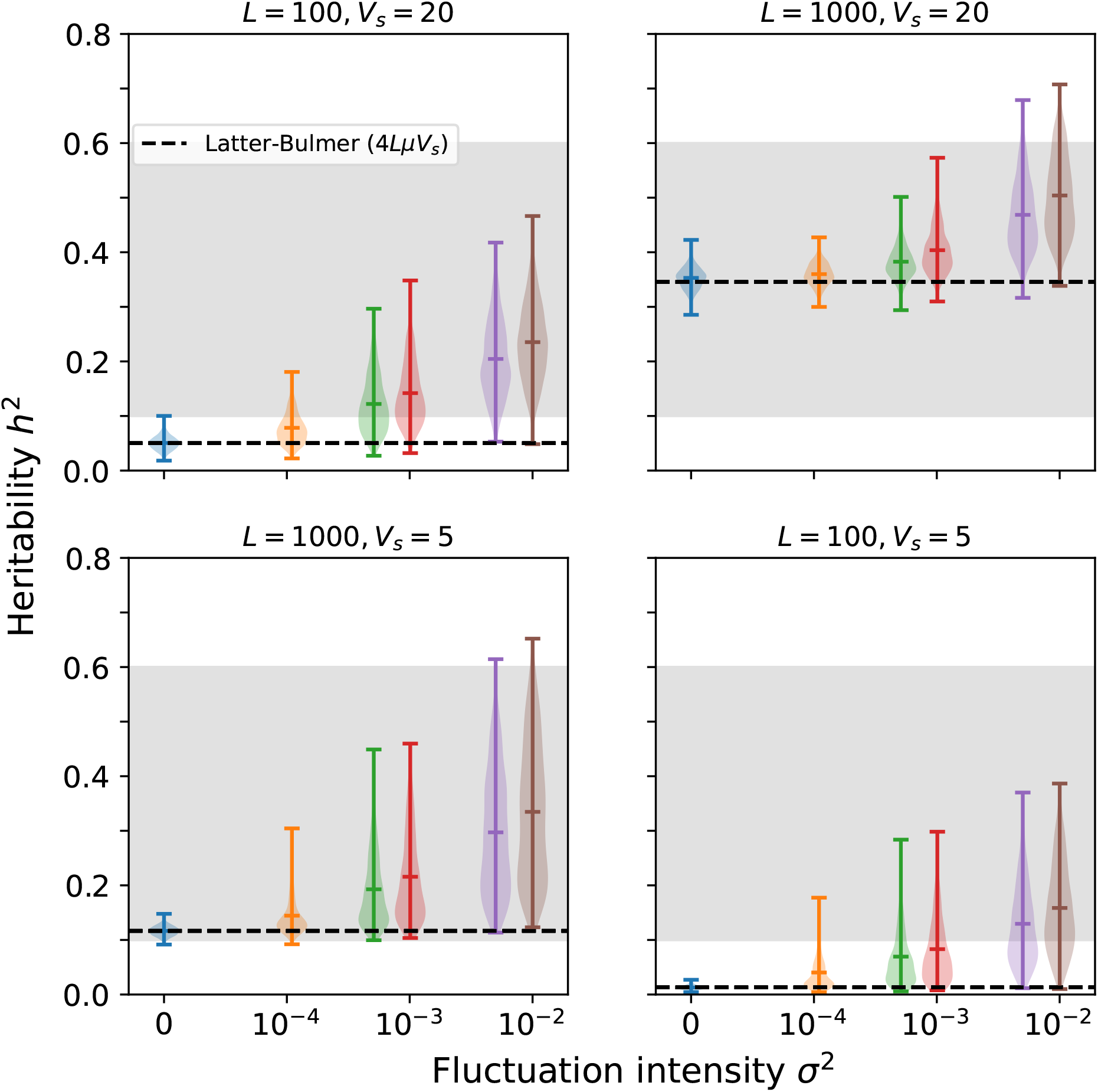
Same as Fig. 2 but with environmental restoring force *θ* = 10^−3^.

**Figure S5:**
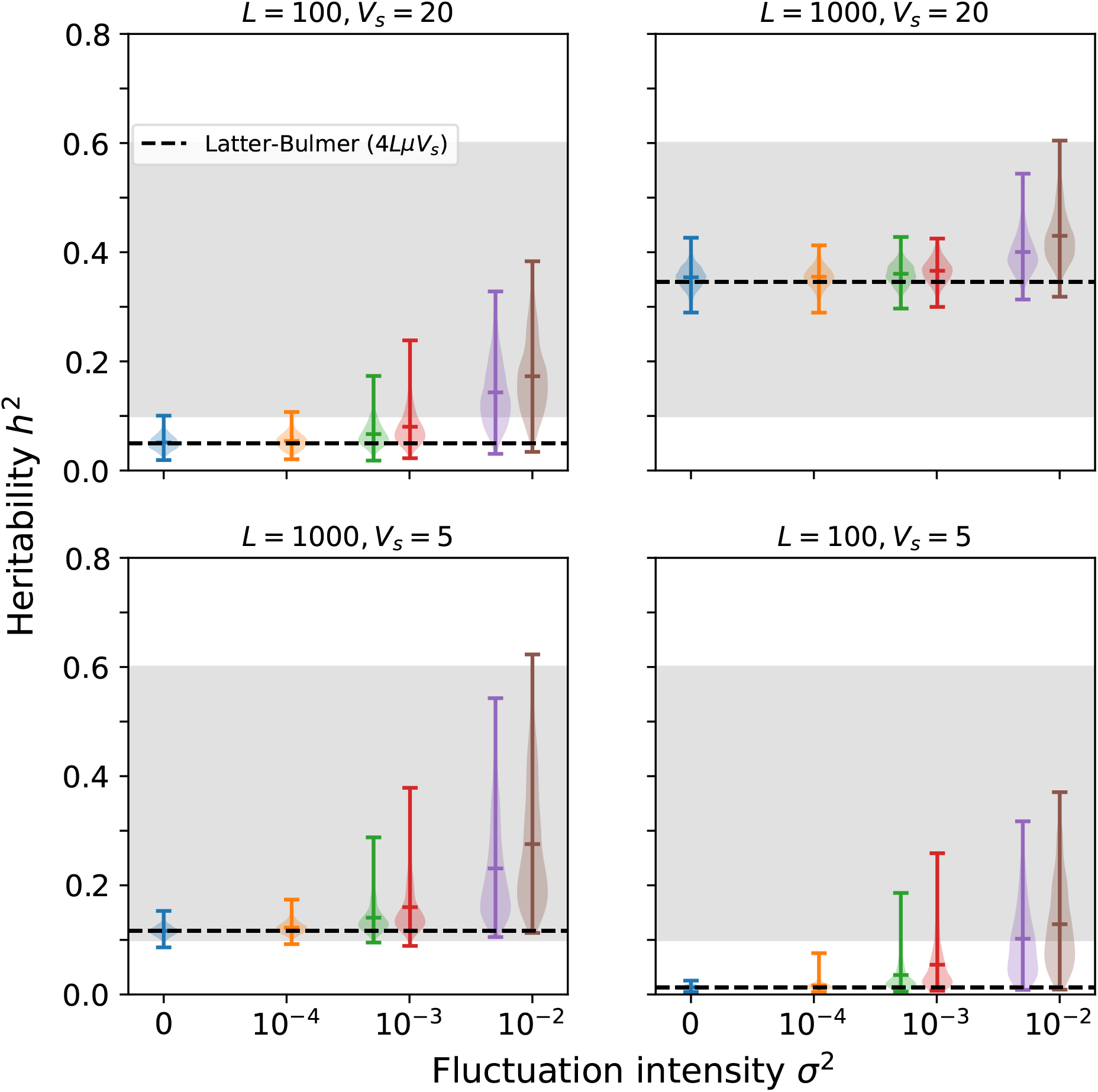
Same as Fig. 2 but with environmental restoring force *θ* = 5×10^−3^.

## References

[1] Neda Barghi, Raymond Tobler, Viola Nolte, Ana Marija Jakšić, François Mallard, Kathrin Anna Otte, Marlies Dolezal, Thomas Taus, Robert Kofler, and Christian Schlötterer. Genetic redundancy fuels polygenic adaptation in drosophila. PLoS biology, 17(2):e3000128, 2019.

[2] Nicholas H Barton. The maintenance of polygenic variation through a balance between mutation and stabilizing selection. Genetics Research, 47(3):209–216, 1986.

[3] Nicholas H Barton. Pleiotropic models of quantitative variation. Genetics, 124(3):773–782, 1990.

[4] Nicholas H Barton and Michael Turelli. Adaptive landscapes, genetic distance and the evolution of quantitative characters. Genetics Research, 49(2):157–173, 1987.

[5] Graham Bell. Fluctuating selection: the perpetual renewal of adaptation in variable environments. Philosophical Transactions of the Royal Society B: Biological Sciences, 365(1537):87–97, 2010.

[6] Jason Bertram and Joanna Masel. Different mechanisms drive the maintenance of polymorphism at loci subject to strong versus weak fluctuating selection. Evolution, 73(5):883–896, 2019.

[7] MG Bulmer. The genetic variability of polygenic characters under optimizing selection, mutation and drift. Genetics Research, 19(1):17–25, 1972.

[8] Reinhard Bürger and Alexander Gimelfarb. Fluctuating environments and the role of mutation in maintaining quantitative genetic variation. Genetics Research, 80(1):31–46, 2002.

[9] Reinhard Bürger and Russell Lande. On the distribution of the mean and variance of a quantitative trait under mutation-selection-drift balance. Genetics, 138(3):901–912, 1994.

[10] Reinhard Bürger and Michael Lynch. Evolution and extinction in a changing environment: a quantitative-genetic analysis. Evolution, 49(1):151–163, 1995.

[11] Reinhard Bürger, Günter P Wagner, and Franz Stettinger. How much heritable variation can be maintained in finite populations by mutation–selection balance? Evolution, 43(8):1748–1766, 1989.

[12] Michael M Desai and Daniel S Fisher. Beneficial mutation–selection balance and the effect of linkage on positive selection. Genetics, 176(3):1759–1798, 2007.

[13] Crispin Gardiner. Stochastic methods, volume 4. Springer Berlin, 2009.

[14] Thomas F Hansen. Stabilizing selection and the comparative analysis of adaptation. Evolution, 51(5):1341–1351, 1997.

[15] Laura Katharine Hayward and Guy Sella. Polygenic adaptation after a sudden change in environment. Elife, 11:e66697, 2022.

[16] Philip W Hedrick. Genetic polymorphism in heterogeneous environments: the age of genomics. Annu. Rev. Ecol. Evol. Syst., 37(1):67–93, 2006.

[17] William G Hill and Mark Kirkpatrick. What animal breeding has taught us about evolution. Annual review of ecology, evolution, and systematics, 41(1):1–19, 2010.

[18] Wen Huang, Richard F Lyman, Rachel A Lyman, Mary Anna Carbone, Susan T Harbison, Michael M Magwire, and Trudy FC Mackay. Spontaneous mutations and the origin and maintenance of quantitative genetic variation. Elife, 5:e14625, 2016.

[19] Emilia Huerta-Sanchez, Rick Durrett, and Carlos D Bustamante. Population genetics of polymorphism and divergence under fluctuating selection. Genetics, 178(1):325–337, 2008.

[20] Kavita Jain and Wolfgang Stephan. Rapid adaptation of a polygenic trait after a sudden environmental shift. Genetics, 206(1):389–406, 2017.

[21] Toby Johnson and Nick Barton. Theoretical models of selection and mutation on quantitative traits. Philosophical Transactions of the Royal Society B: Biological Sciences, 360(1459):1411–1425, 2005.

[22] Samuel Karlin and Benny Levikson. Temporal fluctuations in selection intensities: case of small population size. 1974.

[23] John K Kelly and Kimberly A Hughes. Pervasive linked selection and intermediate-frequency alleles are implicated in an evolve-and-resequencing experiment of drosophila simulans. Genetics, 211(3):943–961, 2019.

[24] Motoo Kimura. Process leading to quasi-fixation of genes in natural populations due to random fluctuation of selection intensities. Genetics, 39(3):280, 1954.

[25] Joel G Kingsolver and Sarah E Diamond. Phenotypic selection in natural populations: what limits directional selection? The American Naturalist, 177(3):346–357, 2011.

[26] Joel G Kingsolver, Hopi E Hoekstra, Jon M Hoekstra, David Berrigan, Sacha N Vignieri, CE Hill, Anhthu Hoang, Patricia Gibert, and Peter Beerli. The strength of phenotypic selection in natural populations. The American Naturalist, 157(3):245–261, 2001.

[27] Alexey S Kondrashov. Rate of evolution in a changing environment. Journal of theoretical biology, 107(2):249–260, 1984.

[28] Alexey S Kondrashov and Lev Yu Yampolsky. High genetic variability under the balance between symmetric mutation and fluctuating stabilizing selection. Genetics Research, 68(2):157–164, 1996.

[29] Russell Lande. Natural selection and random genetic drift in phenotypic evolution. Evolution, pages 314–334, 1976.

[30] BDH Latter. Natural selection for an intermediate optimum. Australian Journal of Biological Sciences, 13(1):30–35, 1960.

[31] Timothy A Mousseau and Derek A Roff. Natural selection and the heritability of fitness components. Heredity, 59(2):181–197, 1987.

[32] Guy Sella and Nicholas H Barton. Thinking about the evolution of complex traits in the era of genome-wide association studies. Annual review of genomics and human genetics, 20:461–493, 2019.

[33] Naoyuki Takahata, Kazushige Ishii, and Hirotsugu Matsuda. Effect of temporal fluctuation of selection coefficient on gene frequency in a population. Proceedings of the National Academy of Sciences, 72(11):4541–4545, 1975.

[34] Naoyuki Takahata and Motoo Kimura. Genetic variability maintained in a finite population under mutation and autocorrelated random fluctuation of selection intensity. Proceedings of the National Academy of Sciences, 76(11):5813–5817, 1979.

[35] Kevin R Thornton. Polygenic adaptation to an environmental shift: temporal dynamics of variation under gaussian stabilizing selection and additive effects on a single trait. Genetics, 213(4):1513–1530, 2019.

[36] Michael Turelli. Heritable genetic variation via mutation-selection balance: Lerch’s zeta meets the abdominal bristle. Theoretical population biology, 25(2):138–193, 1984.

[37] Nicolaas Godfried Van Kampen. Stochastic processes in physics and chemistry, volume 1. Elsevier, 1992.

[38] Bruce Walsh and Mark W Blows. Abundant genetic variation+ strong selection= multivariate genetic constraints: a geometric view of adaptation. Annual review of ecology, evolution, and systematics, 40:41–59, 2009.

[39] Bruce Walsh and Michael Lynch. Evolution and selection of quantitative traits. Oxford University Press, 2018.

[40] Meike J Wittmann, Alan O Bergland, Marcus W Feldman, Paul S Schmidt, and Dmitri A Petrov. Seasonally fluctuating selection can maintain polymorphism at many loci via segregation lift. Proceedings of the National Academy of Sciences, 114(46):E9932–E9941, 2017.

[41] Sewall Wright. The analysis of variance and the correlations between relatives with respect to deviations from an optimum. Journal of Genetics, 30(2):243–256, 1935.

[42] Xiao Yi and Antony M Dean. Bounded population sizes, fluctuating selection and the tempo and mode of coexistence. Proceedings of the National Academy of Sciences, 110(42):16945–16950, 2013.

